# Humans surviving cholera develop antibodies against *Vibrio cholerae* O-specific polysaccharide that inhibit pathogen motility

**DOI:** 10.1101/2020.10.08.332551

**Authors:** Richelle C. Charles, Meagan Kelly, Jenny M. Tam, Aklima Akter, Motaher Hossain, Kamrul Islam, Rajib Biswas, Mohammad Kamruzzaman, Fahima Chowdhury, Ashraful I. Khan, Daniel T. Leung, Ana Weil, Regina C. LaRocque, Taufiqur Rahman Bhuiyan, Atiqur Rahman, Leslie M. Mayo-Smith, Rachel L. Becker, Jatin M. Vyas, Christina S. Faherty, Kourtney P. Nickerson, Samantha Giffen, Alaina S. Ritter, Matthew K. Waldor, Peng Xu, Pavol Kováč, Stephen B. Calderwood, Robert C. Kauffman, Jens Wrammert, Firdausi Qadri, Jason B. Harris, Edward T. Ryan

## Abstract

The mechanism of protection against cholera afforded by previous illness or vaccination is currently unknown. We have recently shown that antibodies targeting O-specific polysaccharide (OSP) of *Vibrio cholerae* correlate highly with protection against cholera. *V. cholerae* is highly motile and possesses a flagellum sheathed in O-specific polysaccharide (OSP), and motility of *V. cholerae* correlates with virulence. Using high speed video microscopy, and building upon previous animal-related work, we demonstrate that sera, polyclonal antibody fractions, and OSP-specific monoclonal antibodies recovered from humans surviving cholera block *V. cholerae* motility at both subagglutinating and agglutinating concentrations. This anti-motility effect is reversed by pre-adsorbing sera and polyclonal antibody fractions with purified OSP; and is associated with OSP-specific but not flagellin-specific monoclonal antibodies. F[ab] fragments of OSP-specific polyclonal antibodies do not inhibit motility, suggesting a requirement for antibody-mediated crosslinking in motility inhibition. We show that OSP-specific antibodies do not directly affect *V. cholerae* viability, but that OSP-specific monoclonal antibody highly protects against death in the murine cholera model. We used *in vivo* competitive index studies to demonstrate that OSP-specific antibodies impede colonization and survival of *V. cholerae* in intestinal tissues, and that this impact is motility-dependent. Our findings suggest that the impedance of motility by antibodies targeting *V. cholerae* OSP contributes to protection against cholera.

**IMPORTANCE:** Cholera is a severe dehydrating illness of humans caused by *Vibrio cholerae. V. cholerae* is a highly motile bacterium that has a single flagellum covered in lipopolysaccharide (LPS) displaying O-specific polysaccharide (OSP), and *V. cholerae* motility correlates with its ability to cause disease. The mechanisms of protection against cholera are not well understood; however, since *V. cholerae* is a non-invasive intestinal pathogen, it is likely that antibodies that bind the pathogen or its products in the intestinal lumen contribute to protection from infection. Here, we demonstrate that OSP-specific antibodies isolated from humans surviving cholera in Bangladesh inhibit *V. cholerae* motility and are associated with protection against challenge in a motility-dependent manner.

## INTRODUCTION

Cholera is a severe dehydrating illness of humans caused almost exclusively by *Vibrio cholerae* of the O1 serogroup. Over one billion people remain at risk for cholera in 51 endemic countries, and there are an estimated 3 million cases and 95,000 deaths each year from cholera (1). The current global pandemic began in 1961 and gives no indication of abating, as evidenced by recent large outbreaks in Haiti and Yemen (2). This reality has led to enhanced commitments to cholera control strategies (3). Such strategies now include vaccination against cholera, along with efforts to improve water and sanitation (4). Currently available oral killed cholera vaccines are an important addition to these control efforts; however, these vaccines may provide limited durable protection, especially in immunologically naïve individuals, including children under 5 years of age who bear a large proportion of the global cholera burden (2). In comparison, survivors of cholera, including young children, have high level protective immunity that persists for years (5).

The development and optimal use of cholera vaccines has been hampered by the relatively limited understanding of the immunologic mechanisms of protection against cholera. *V. cholerae* is a non-invasive luminal intestinal pathogen, and it is likely that antibodies that bind the pathogen or its products in the intestinal lumen contribute to protective immunity (6). Cholera is a toxin-mediated disease; *V. cholerae* express cholera toxin (CT), an ADP-ribosylating enzyme, at the intestinal surface, and the actions of this toxin on intestinal epithelial cells lead to the large volume secretory diarrhea characteristic of cholera (7). Despite this, immune responses that target CT do not provide meaningful protection against cholera (8).

An *in vitro* vibriocidal assay is currently our best predictor of protection against cholera; however, the vibriocidal response appears to be a surrogate marker of an as yet to be identified mucosal antibody response(s) (7). We have shown that the vibriocidal response largely targets the O-specific polysaccharide (OSP) of *V. cholerae* (9). Moreover, we found that OSP-specific antibody and memory B cell responses correlate with protection against cholera in household contacts of cholera index patients in Bangladesh (10). In North American recipients of an oral cholera vaccine, OSP-specific antibody responses correlate with protection against cholera in challenge studies (11).

How OSP-specific antibodies protect against *V. cholerae* in the intestinal lumen is currently unclear. Possible mechanisms include direct bacteriocidal, enchaining, or agglutinating activity (12). However, we hypothesized that inhibition of motility could be a potential mechanism as well; a possibility supported by previous work (13-16). *V. cholerae* is a highly motile bacterium that has a single polar flagellum sheathed in lipopolysaccharide (LPS) displaying OSP, and *V. cholerae* motility correlates with virulence (17-19). Furthermore, several *in vitro* studies have shown that antibodies targeting *V. cholerae* LPS impede *V. cholerae* motility (13-16), and studies in suckling mice have suggested that the impedance of motility by anti-OSP antibodies provides protection in this model (13, 14, 16, 20, 21). Here, we used antibodies recovered from humans surviving cholera in Bangladesh, including monoclonal antibodies cloned from plasmablasts homing to the intestinal mucosa (22), to investigate the role of anti-OSP antibodies in protection against cholera. Our findings suggest that the inhibition of motility by the bivalent binding of anti-OSP antibodies contributes to immune protection from cholera.

## RESULTS

### Polyclonal OSP-specific antibodies inhibit *V. cholerae* motility

We analyzed polyclonal antibody responses in plasma from ten adults recovering from cholera caused by *V. cholerae* O1 Ogawa in Bangladesh (Table S1). These patients all developed IgG, IgA and IgM responses against *V. cholerae* O1 OSP (p<0.01 for all antibody isotypes; Fig. 1A). The anti-*V. cholerae* O1 Ogawa agglutination activity was present in plasma across a range of dilutions in individual patients (1:64 – >1:8) (Table 1). Using high speed video dark-field microscopy, we then assessed motility after incubating *V. cholerae* O1 with acute and convalescent phase heat-inactivated plasma in sub-agglutinating concentrations (1:256) in the same conditions used for the agglutination assays. There was a significant decrease in motility when *V. cholerae* organisms were mixed with convalescent plasma compared to acute plasma (p = 0.001; Fig. 1B). Adsorbing plasma with purified *V. cholerae* OSP eliminated its capacity to inhibit *V. cholerae* motility (Fig. 1B).

**Table 1:**
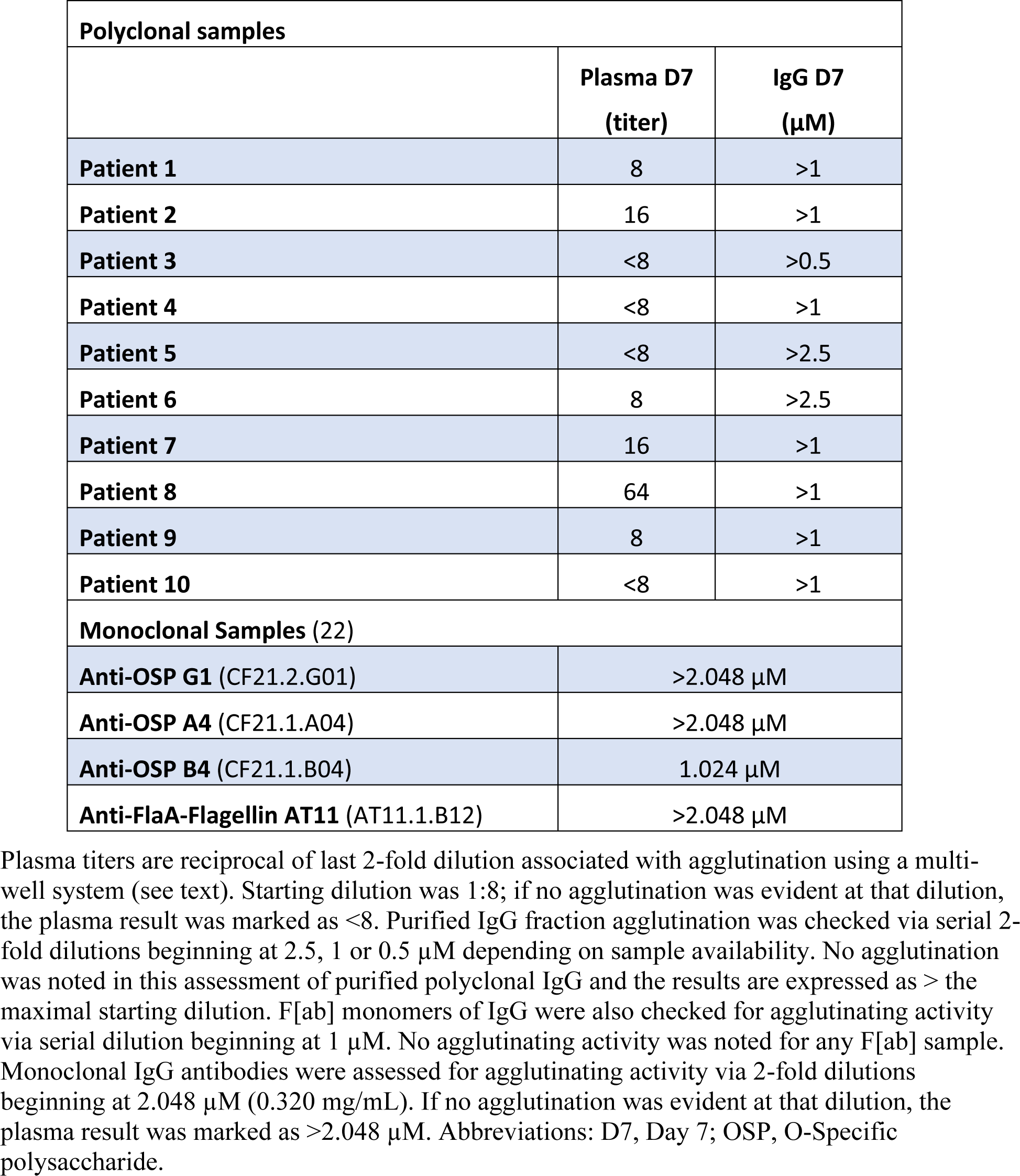
*V. cholerae* O1 Ogawa agglutination activity of antibody samples.

**Fig. 1.**
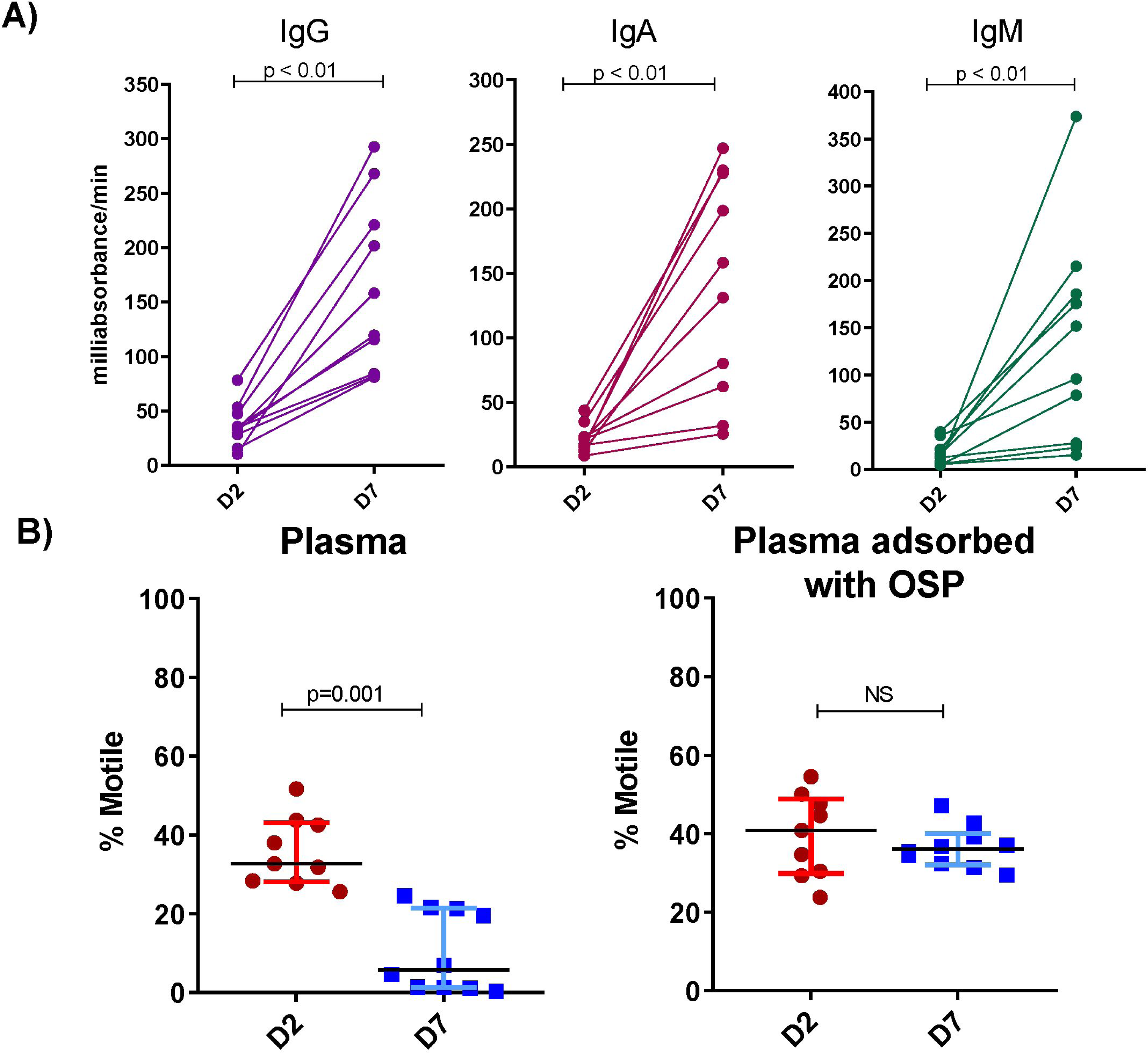
Convalescent phase plasma of cholera patients recognizes *V. cholerae* OSP and inhibits *V. cholerae* motility in an OSP-dependent manner. **(A)** IgG, IgA, and IgM plasma responses targeting *V. cholerae* O1 Ogawa OSP at acute (D2) and convalescent (D7) phase of cholera (n=10) as determined by enzyme-linked immunosorbent assay. **(B)** Motility of *V. cholerae* O1 Ogawa O395 assessed by high speed video microscopy after a 5-minute incubation with a sub-agglutinating dilution of plasma (1:256) of cholera patients (n=10) at a bacterial OD_600_ of 0.1.. Percentages of motile versus non-motile bacteria after incubation with heat inactivated-plasma with and without adsorption with *V. cholerae* OSP are shown. Error bars = median with interquartile range. Differences within groups were assessed using the Wilcoxon matched-pairs signed rank sums test. Abbreviations: D2, day 2; D7, day 7; OSP, O-specific polysaccharide.

We next isolated IgG, IgA and IgM antibody fractions from these plasma samples, and focused our analyses on the IgG fraction, the most abundant isotype recovered. Using conditions that replicated those used in the video microscopy, we did not detect agglutination in 2-fold dilutions of purified IgG starting at 0.5, 1 or 2.5 μM depending on sample availability (Table 1). Starting at a concentration of 2.5 μM, we found the impact of pooled IgG on *V. cholerae* motility was concentration dependent and detectable at a subagglutinating concentration of 0.25 μM (Fig. 2A; p ≤0.05) and independent of effects on *V. cholerae* viability (Fig. S1). Purified IgG fractions from individual patients had similar anti-OSP reactivity (Fig 2B; p ≤0.01), and inhibition of *V. cholerae* motility at 0.25 μM was abrogated by adsorbing the IgG with purified *V. cholerae* OSP (Fig. 2C; p ≤0.05). The absence of bacterial clumping/agglutination at 0.25 μM IgG was confirmed by microscopy. Thus, polyclonal IgG anti-OSP antibodies in convalescent patient sera inhibit the cholera pathogen.s motility at concentrations that do not cause the organism to agglutinate.

**Fig. 2:**
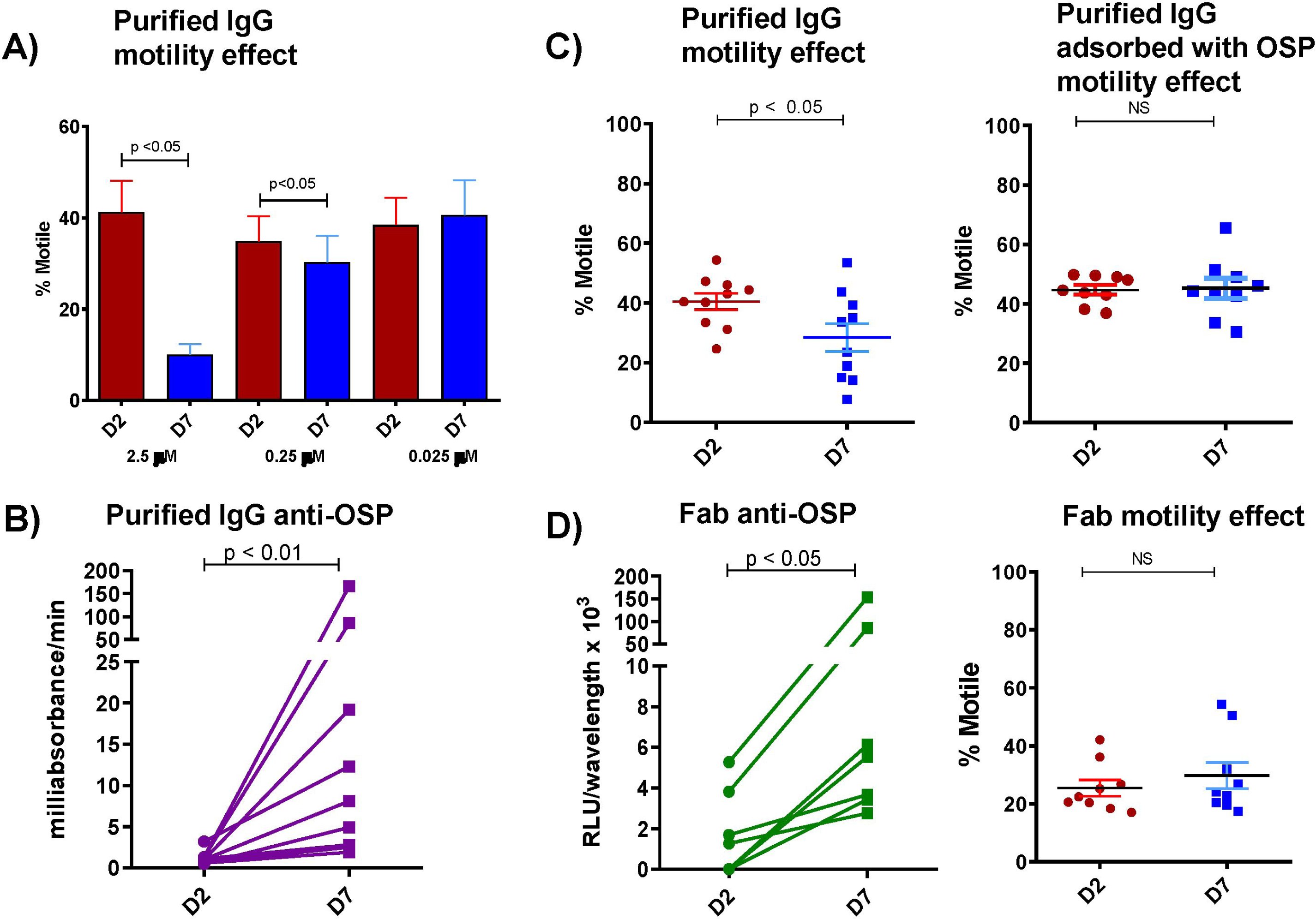
Purified IgG from humans recovering from cholera inhibits *V. cholerae* motility in a concentration dependent manner and this effect is eliminated when IgG is adsorbed with *V. cholerae* OSP. (**A**) IgG purified from pooled convalescent phase plasma of humans recovering from cholera inhibits *V. cholerae* motility in a concentration dependent manner. (**B**) OSP-specific responses of purified IgG from individual patients measured by enzyme-linked immunofluorescence assay showing day 7 immunoreactivity. (**C**) Purified polyclonal IgG at a sub-agglutinating 0.25 μM concentration from humans recovering from cholera inhibits *V. cholerae* motility and this effect is eliminated when IgG is adsorbed with *V. cholerae* OSP. (**D**) F[ab] fragments generated from purified IgG retain day 7 immunoreactivity to OSP but lose ability to inhibit *V. cholerae* motility. D2: Day 2; D7: Day 7. Differences within groups were assessed using the Wilcoxon matched-pairs signed rank sums test.

To assess whether antibody-based cross-linking was required for inhibition of *V. cholerae* motility, we generated Fragment antigen-binding (F[ab]) fragments from the IgG fractions of individual patients. In contrast to intact IgG antibodies, which both bound *V. cholerae* OSP (Fig 2B; p <0.01) and blocked *V. cholerae* motility (Fig. 2C; p ≤0.05), F[ab] fragments bound OSP (Fig. 2D; p ≤0.01), but did not inhibit motility (Fig. 2D), suggesting that multivalent binding of OSP by antibody molecules is important for impeding motility. Supporting this idea, we found that dimeric IgA and polymeric IgM fractions also inhibited *V. cholerae* motility in an OSP-dependent fashion (Fig. S2; p ≤0.001).

### Monoclonal OSP-specific antibodies inhibit *V. cholerae* motility

We next turned our attention to analyses of *V. cholerae*-specific monoclonal antibodies isolated from intestinal mucosal-homing plasmablasts from the peripheral blood of cholera patients sampled 7 days after they presented for clinical care in Bangladesh (22). We previously cloned (as human IgG1) and characterized over 100 monoclonal antibodies from this collection (22). For the current analysis, we selected 4 previously characterized antibodies including one monoclonal antibody with high affinity for *V. cholerae* OSP (G1), two antibodies with low affinity for *V. cholerae* OSP (A4, B4), and one specific for *V. cholerae* flagellin A (AT11) (22). The OSP-specific monoclonal IgG antibodies were previously shown to recognize both Ogawa and Inaba antigens(22). Monoclonal IgG antibodies were assessed for agglutinating activity via 2-fold dilutions beginning at 2.048 μM (0.320 mg/mL). If no agglutination was evident at this dilution, the plasma result was marked as >2.048 μM. One low affinity OSP-specific IgG monoclonal antibody (B4) had agglutinating activity at 1.024 μM, whereas the other three monoclonal antibodies (G1, A4, and AT11) did not have detectable agglutinating activity at the highest concentration assessed (2.048 μM) (Table 1). Similar to our findings using polyclonal antibodies described above, we found that the anti-OSP monoclonal antibodies also blocked *V. cholerae* motility in a concentration dependent manner, including at sub-agglutinating concentrations, whereas the anti-flagellin antibody did not block motility (Fig. 3; p ≤0.001-0.05; Movies S1, S3). Inhibition of ‘shooting star’ motility at the single bacterial cell level (without agglutination and clumping) was demonstrated by analysis of freeze-frame images of *V. cholerae* in the presence of human monoclonal OSP-specific antibodies, but not in the presence of the anti-flagellin antibody (Fig. 3, 4; Movies S1, S2, S3). There were no apparent differences in the capacities of the high versus low affinity OSP-specific antibodies to inhibit *V. cholerae* motility (Fig. 3,4; Movies S1, S2). Scanning electron microscope imaging of *V. cholerae* revealed that OSP-specific monoclonal antibody promoted flagellar tethering, but tethering was occasionally observed in the presence of flagellar-specific monoclonal antibody (Fig. S3). Occasional bacterial blebbing was also observed in scanning electron microscopic analyses with both antibodies. Our attempts to confirm the potential requirement for antibody mediated cross-linking using the monoclonal antibodies were confounded by aggregation of monoclonal F[ab] fragments in all assessed non-reducing conditions.

**Fig. 3.**
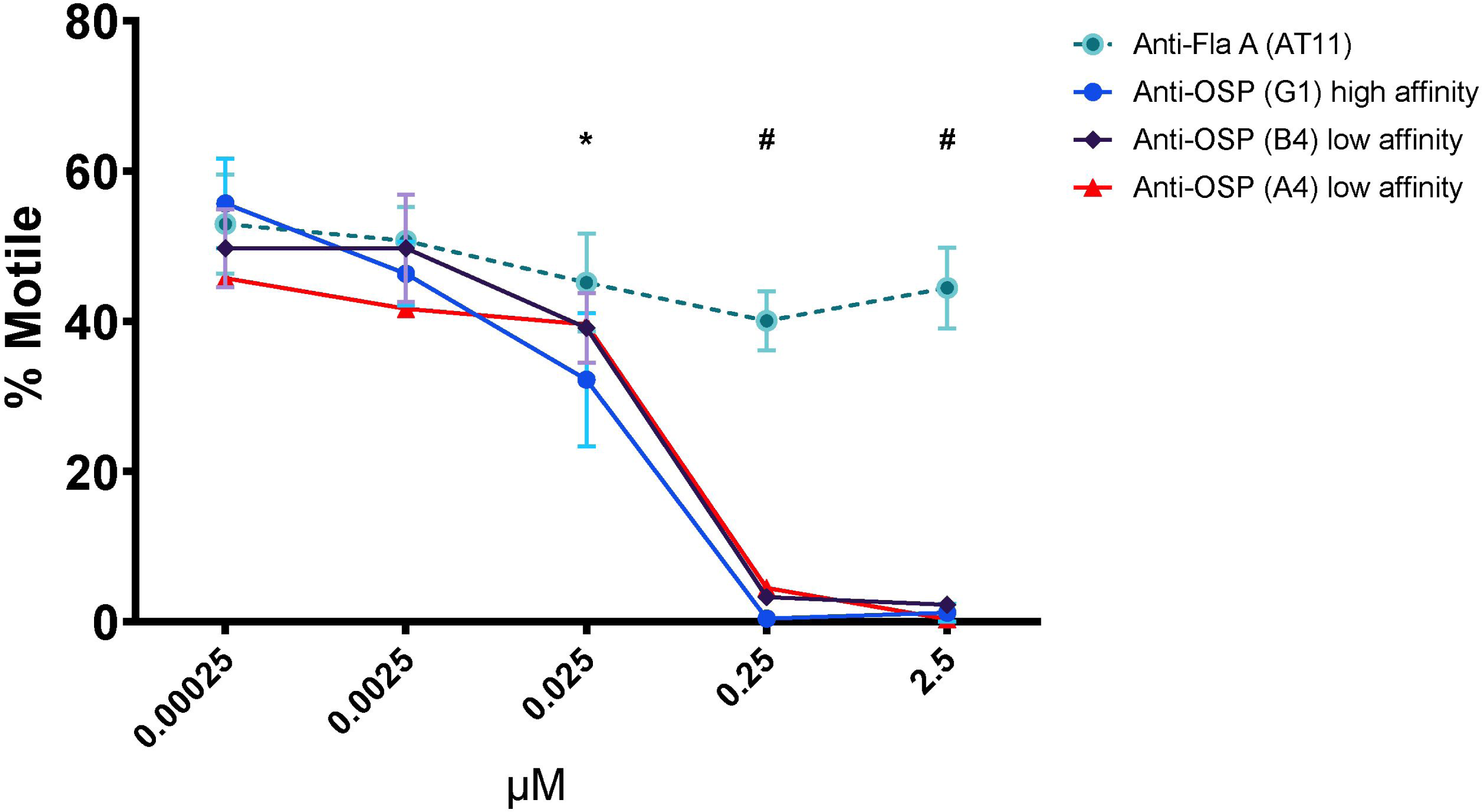
Human monoclonal antibodies targeting *V. cholerae* OSP inhibit motility in a concentration dependent manner. Motility of *V. cholerae* O1 (OD_600_=0.1) was measured by phase contrast video microscopy after a 5-minute incubation with increasing concentrations of human monoclonal IgG antibodies targeting OSP (high and low affinity) or Flagellin A (FlaA). Each point represents the mean of experiments performed in triplicate. Error bars = standard error of the mean. * p ≤0.05 G1 compared to 0.00025 μM; # p ≤0.001 G1, A4, B4 compared to 0.00025 μM.

**Fig. 4.** Freeze-frame phase contrast demonstrating anti-OSP human monoclonal antibodies eliminate *V. cholerae* shooting star motility, while anti-flagellin monoclonal antibody does not. Freeze frame (400ms) images of *V. cholerae* O1 Ogawa O395 (OD 0.1) after 5-minute incubation with phosphate-buffered saline, or with human monoclonal antibodies (0.25 μM) targeting OSP (G1, B4) or FlaA. Lines represent motile bacteria (shooting star); white dots represent non-motile bacteria. Images were taken from high speed videos and are representative of experiments performed in triplicate. Note elimination of white streaks demonstrating shooting star *V. cholerae* motility in the presence of OSP-specific monoclonal antibodies.

### OSP-specific monoclonal antibody promotes survival and inhibits *V. cholerae* intestinal colonization in mice

We used a neonatal mouse challenge model to determine whether OSP-specific monoclonal antibody G1 alters the survival of suckling mice after lethal challenge with wild type *V. cholerae* O1 strain C6706. Mixture of G1 with C6706 prior to oro-gastric inoculation led to a marked reduction in death of the mice (Fig. 5A; p < 0.001 value). To assess how the effect of OSP-specific antibody on motility impacts *V. cholerae* survival and localization in the intestine, we carried out competition studies where the colonization of *V. cholerae* C6706*lacZ*^-^ (a strain that colonizes as well as the wild type) was compared to either 1) a C6706*lacZ*^+^ transposon mutant that was flagellated but non-motile (*motB*::Kan^r^) or to 2) a C6706*lacZ*^+^ transposon mutant that was motile but rough and lacking OSP due to deficient perosamine synthesis (*VC0244*::Kan^r^). The motilities of the comparator and rough strains were equivalent, whereas the *motB*::Kan^r^ strain was confirmed as non-motile. Relative to the C6706*lacZ*^-^ strain, the non-motile *motB*::Kan^r^ mutant had a marked thousand-fold colonization defect in the proximal small intestine and a less pronounced 100-fold defect in the distal small intestine (Fig. 5B). When an OSP-specific monoclonal antibody was added to the inocula mixture at subagglutinating concentration, the competitive disadvantage of the non-motile mutant was significantly ameliorated in the proximal but not distal small intestine (Fig. 5B; p <0.001). This reduction in the competitive defect related to a decrease in recovery of motile *V. cholerae* from the proximal small intestine in the presence of OSP-specific antibody G1 (Table S3). In contrast, addition of the anti-flagellar monoclonal antibody to the inocula did not alter the competitive advantage of motile *V. cholerae* compared to the non-motile mutant in either the proximal or distal intestine (Fig. 5B). These observations suggest that the inhibition of motility by anti-OSP antibody impedes *V. cholerae*.s capacity to colonize the proximal small intestine.

**Fig. 5:**
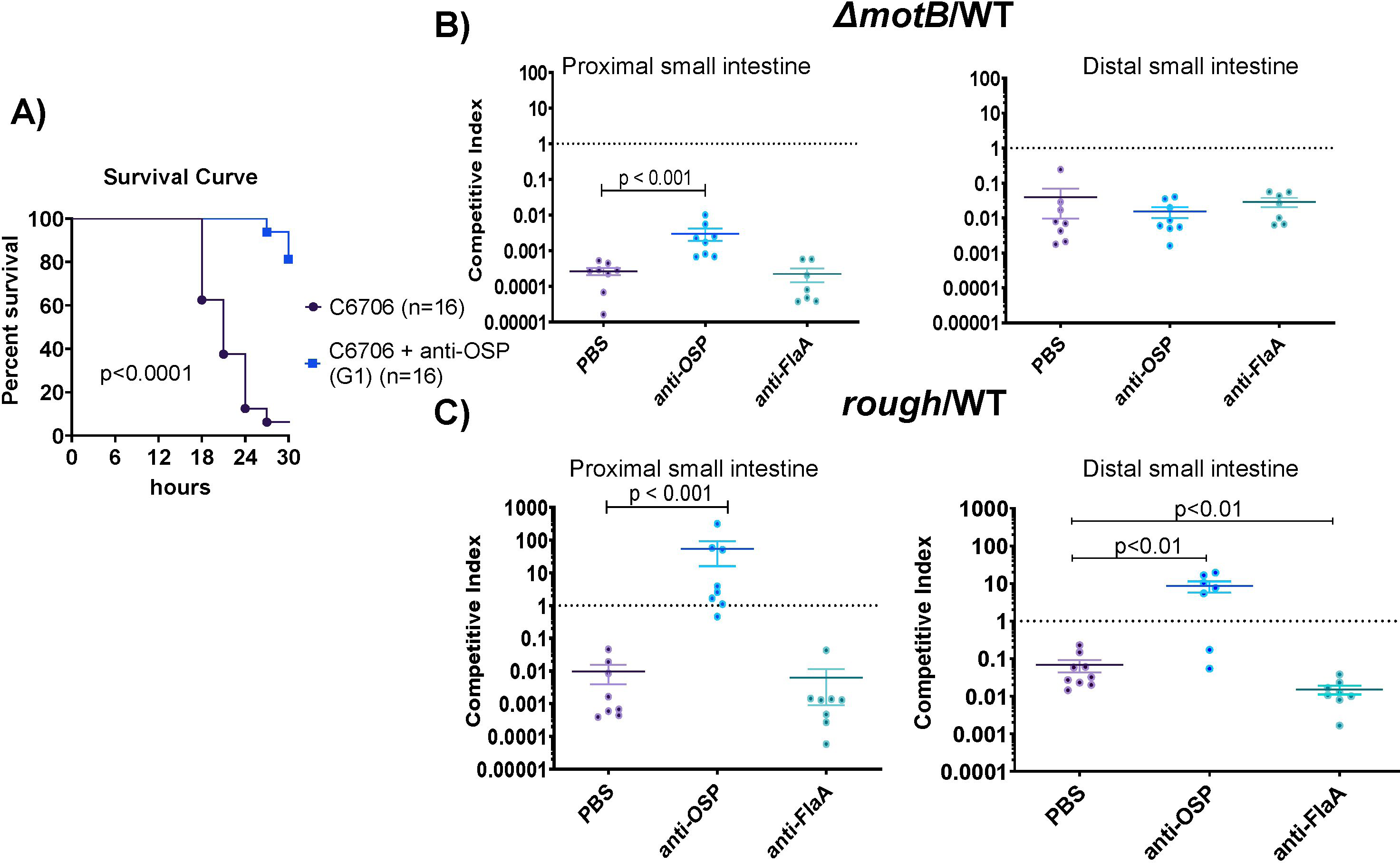
Impact of OSP-specific monoclonal antibodies on *V. cholerae* lethality and intestinal colonization in suckling mice. **(A)** OSP monoclonal antibody protects against death in the mouse neonatal *V. cholerae* challenge model. Wild type *V. cholerae* O1 strain C6706 (10^9^ organisms) was mixed with 0.25 μM human monoclonal antibody targeting *V. cholerae* OSP (G1) or PBS. Fifty μl of sample mixture was inoculated into 3 to 5-day old sucking CD-1 mice and survival was assessed over a 30-hour period every 3 hours. Survival curves were compared by log rank testing. PBS, phosphate buffered saline. **(B, C)** Inhibition of *V. cholerae* motility by OSP-specific monoclonal antibody prevents colonization of the proximal small intestine. *V. cholerae* strain C6706*lacZ*^-^ (WT*LacZ*^-^) was competed against transposon mutants: (**B**) non-motile (*motB*::Kan^r^) or (**C**) rough strains (*VC0244*::Kan^r^) in 3-5 day old suckling mice with the addition of PBS, human mAB targeting *V. cholerae* OSP (G1), or Flagellin A (FlaA). The competitive index (CI) was calculated for the proximal third and distal third of the small intestine as the output ratio of mutant to WT*lacZ*^-^ strain divided by the input ratio of mutant to WT*lacZ*^-^. Each symbol represents CI from a single mouse. Error bars represent the standard error of the mean. Differences between groups were assessed using a Mann-Whitney test. Abbreviations: PBS – phosphate-buffered saline; OSP, O-specific polysaccharide; FlaA, Flagellin A.

In similar experiments where the comparator smooth C6706*lacZ*^-^ strain was competed with the rough (OSP^-^) mutant, we found that the rough strain had a ∼100-fold colonization defect in both the proximal and distal small intestine (Fig, 5C). Notably, recovery of the smooth comparator strain decreased in the presence of OSP-specific monoclonal antibody, in part abolishing the disadvantage of the rough strain in both proximal and distal small intestine (Fig. 5C; p<0.001; Table S3). In marked contrast, the anti-flagella monoclonal antibody did not alter the competitive indices found in samples from intestinal samples.

## DISCUSSION

We found that adult humans recovering from cholera develop antibody responses that inhibit *V. cholerae* motility. These antibodies, which impact motility at subagglutinating concentrations of antibody, target *V. cholerae* OSP and may mediate their effect via cross-linking. This work extends previous animal and *in vitro* LPS-based analyses (13-16).

*V. cholerae* motility is associated with pathogenesis (23). *V. cholerae* have a single sheathed polar flagellum, and swim rapidly in an arcing pattern that can be visualized on dark field microscopy of freshly passed cholera stools. This *V. cholerae* distinctive swimming pattern can be used to rapidly identify *V. cholerae* in the stool of potentially infected patients via dark field microscopy and is referred to as “shooting star” motility (24). As shown in our videos (Movies S1-3), the impact of anti-OSP antibody on *V. cholerae* “shooting star” motility is significant. Motility is involved not only with horizontal movement of *V. cholerae* in the intestine (along the axis from stomach to colon), but also vertical localization in intestinal tissue (most importantly, along the axis created by villi and crypts in the small intestine) (25). Once ingested, the few *V. cholerae* that survive the acidic environment of the stomach enter the small intestine, and a subset swim into the lower villi and intestinal crypts (25). Motility is required for this vertical localization (25). *V. cholerae* motility is then involved in initial penetration of intestinal mucin that overlays the intestinal epithelium (26). Following this initial penetration, *V. cholerae* become non-motile (with many bacteria losing their flagella), transit the mucin (in a poorly understood process), adhere at the intestinal epithelial surface, where they form microcolonies, and secrete cholera toxin (26). This transition from a motile bacterium to an adherent microaggregated form at the intestinal surface is closely controlled by an interplay of quorum sensing by LuxO and HapR, motility, and *V. cholerae* virulence networks (26-33).

Following microcolony formation and the actions of cholera toxin on the intestinal epithelial cell leading to fluid secretion into the intestinal lumen, microcolonies break apart and *V. cholerae* regain their motility in a poorly understood process. This “mucosal escape response” involves RpoS-dependent down regulation of virulence genes, and upregulation of motility genes (34). *V. cholerae* motility facilitates release of organisms from microcolonies (35). Detached organisms are then flushed/swim into the intestinal lumen and are passed into the environment in the rice water stools characteristic of cholera. These highly motile organisms have high transmission potential, a state that has been termed the “hyper-infectious state” (36, 37).

Based on this sequence of events, inhibition of *V. cholerae* mobility could impact *V. cholerae* pathogenesis at a number of critical points. Inhibition of motility could alter intestinal passage, reducing the number of bacteria that move horizontally through the intestinal tract. It could also impact the ability of *V. cholerae* to reach optimal ecologic niches in the lower third of intestinal villi and crypts and could affect the ability of *V. cholerae* to penetrate the protective mucin layer overlying the intestinal epithelium. Here, using the suckling mouse model, we found that the largest impact of anti-OSP antibodies was in the proximal small intestine, the region with the thickest mucin layer (25). Given the prominent interactions among the *V. cholerae* quorum sensing, motility and virulence networks, the inhibition of motility by anti-OSP antibodies could modify the pathogen.s intestinal colonization, toxin expression, and virulence. Altered *V. cholerae* motility could also affect the ability of microcolonies to break apart and for *V. cholerae* to re-enter the intestinal lumen and be flushed from the patient, thus affecting the hyper-infectious and transmission potential of passed *V. cholerae*.

Our findings suggest that the effect of OSP-specific antibodies on *V. cholerae* motility requires at least bivalent binding of antibody. F[ab] fragments of polyclonal antibodies retained their capacity to bind OSP but did not impede *V. cholerae* motility. We also found that convalescent phase IgM and IgA antibodies had a greater effect on *V. cholerae* motility than IgG; this may be secondary to the pentavalent and dimeric structures of IgM and IgA, respectively, which markedly increase their valency. Other groups have assessed the impact of murine antibodies on *V. cholerae* motility with variable results. Analysis of F[ab] fragments generated from polyclonal antibody generated in mice vaccinated against *V. cholerae* suggested that elimination of multi-valency eliminated the effect of the antibodies on *V. cholerae* motility (16). However, F[ab] fragments generated from a number of murine monoclonal anti-LPS antibody apparently affect *V. cholerae* motility (14, 15, 38).

Our observations may help explain this discrepancy. Despite clearly demonstrating complete cleavage of antibody and removal of the Fc (Fragment, crystallizable) fraction, we were unable to prevent aggregation of F[ab] fragments of human monoclonal antibodies in any physiologic non-reducing condition at relevant concentrations. We did not observe aggregation in F[ab] fragments generated from human polyclonal antibody, which were used in our analyses. The propensity of fragmented monoclonal antibodies to form aggregates, presumably due to their inability to form proper disulfide bonds, has been described previously (39).

The requirement for multivalent binding to affect *V. cholerae* motility suggests at least two possible mechanisms for OSP-specific effect on *V. cholerae* motility, depending on the concentration of bacteria and antibody and length of interaction time. The first requires sufficient bacteria, antibody and time and involves interbacterial cross-linking of bacteria, *i*.*e*. agglutination. The second involves only the concentration of antibody and impact on individual bacteria. Our results strongly suggest that both conditions can impact *V. cholerae* motility, and in the human intestine, both conditions could exist. Not surprisingly, we were able to show an impact on *V. cholerae* motility in agglutinating conditions. More interestingly, we demonstrated the impact of human anti-OSP antibodies at antibody concentrations less than those associated with agglutination, and we were able to confirm inhibition of motility at the single bacterial level by microscopy. How anti-OSP antibodies could affect *V. cholerae* motility at subagglutinating concentrations is currently unclear. One possibility is flagellar tethering or kinking due to antibody-mediated OSP cross-linking (14). We did indeed detect flagellar tethering using electron microscopy, although we also observed occasional tethering with control antibody as well. Other mechanisms of action are also possible. For instance, the colonization defect of the flagellated but non-motile *motB* mutant strain in comparison to the wild type strain was not completely reversed when both were exposed to anti-OSP antibody, suggesting that the genetically non-motile strain may be less able to respond to intestinal environmental signals and activate colonization and virulence cascades (40). An effect on transcriptional or protein expression in the setting of OSP-specific binding is also possible (41).

A limitation of our study was our inability to assess IgA and IgM fractions due to the low amounts recovered from polyclonal serum. *V. cholerae* is a non-invasive mucosal infection, and IgM and IgA are thought to be the predominant antibodies involved in mediating protection (11). We were, however, able to include analysis of monoclonal antibodies cloned from mucosal-homing plasmablasts in patients with cholera (22), and analyses comparing the impacts of IgM, IgA and IgG isotype anti-OSP antibodies on *V. cholerae* motility are planned.

Despite this limitation, our results show that humans surviving cholera develop antibodies that inhibit *V. cholerae* motility through binding of the OSP component of LPS, at both subagglutinating as well as agglutinating conditions. These data build upon a growing body of evidence that OSP-specific antibody responses mediate protection against cholera and suggest that such antibodies may act by mechanisms other than agglutination. Finally, our findings also support a novel mechanism of antibody-mediated protection through direct inhibition of bacterial motility.

## MATERIALS AND METHODS

### Study subject selection and sample collection

Venous blood was collected at the acute phase (day 2) and convalescent phase of infection (day 7) from ten adult patients (age 18-55 years) presenting to the International Centre for Diarrhoeal Disease Research, Bangladesh (icddr,b) hospital with culture-confirmed *V. cholerae* O1 El Tor Ogawa infection. Patients were treated with intravenous fluids and/or oral rehydration solution and antimicrobials at the discretion of the attending physician. This study was approved by the Research and Ethical Review Committees of the (icddr,b) and the human studies committee of Massachusetts General Hospital.

### Bacterial strains

The *V. cholerae* O1 strains used in this study are listed in Table S2 (42). Bacterial strains were grown at 37 °C in Luria-Bertani (LB) media with or without antibiotics as indicated. Antibiotic concentrations were: streptomycin (Sm) at concentration of 100 μg/mL and kanamycin (Km) 45 μg/mL. X-gal (200 μg/ml) plus isopropyl β-D-1-thiogalactopyranoside (IPTG; 0.1 mM) was used for blue/white colony screening. For experiments with the rough strain, bacteria were resuspended in 0.5% NaCl to prevent autoagglutination after growth in LB.

### Purification of antibody isotypes from plasma

IgG was purified from heat-inactivated human plasma using Protein G HP spin trap columns (GE Healthcare) and then incubated with CaptureSelect IgA Affinity Matrix (ThermoScientific) to remove contaminating IgA per manufacturer.s instructions. IgA and IgM antibodies were then subsequently purified by sequentially running the flow through from the Protein G column onto columns loaded with CaptureSelect IgA and IgM Affinity Matrix (ThermoScientific). Quantity and purity of IgG, IgA, and IgM isotype fractions was determined by ELISA using standard curves of ChromPure human IgG, IgA, and IgM (Jackson ImmunoResearch).

### F[ab] fragmentation of plasma IgG

F[ab] fragments of IgG were generated using the F[ab] Preparation Kit (Pierce) per the manufacturer.s instructions. Briefly, immobilized papain was used to digest the desalted and purified IgG, and the resultant F[ab] fragments were purified using a Protein A Plus spin kit (Pierce). Purity of the F[ab] fragments was assessed by Western blot analysis with 100 ng of sample per well and direct probing with anti-IgG F[ab]_2_ and anti-IgG Fc conjugated to horseradish peroxidase (Jackson ImmunoResearch).

### Monoclonal antibody production

Recombinant human monoclonals were generated from the cholera-induced day 7 plasmablast population of *V. cholerae*-infected Bangladeshi patients by single-cell expression as previously described(22). For this analysis, we used the following previously characterized human IgG monoclonal antibodies: high affinity anti-OSP G1 (CF21.2.G01); low affinity anti-OSP A4 (CF21.1.A04); low affinity anti-OSP B4 (CF21.1.B04); anti-FlaA-Flagellin AT11 (AT11.1.B12) (22).

### Generation of *V. cholerae* O1 OSP

OSP from *V. cholerae* O1 Ogawa strain PIC158 was purified and conjugated to BSA as previously described (43).

### Vibriocidal assays

Vibriocidal assays were performed on heat-inactivated plasma samples as previously described (44) and used target strain *V. cholerae* O1 Ogawa strain O395. Heat-inactivated plasma and exogenous guinea pig complement (EMD Millipore) were incubated with the target strain. Vibriocidal titers were defined as the reciprocal of the highest serum dilution resulting in a 50% reduction in optical density (OD_600_) at 595 nm absorbance compared to no serum controls.

### OSP ELISA

OSP responses were measured using ELISA as previously described (44). Microplates were coated with 100 ng/well of O1 Ogawa OSP:BSA. Samples were added to plates (plasma 1:50, purified IgG 10 μM; purified F[ab] 1 μM), and IgG responses were detected with goat anti-human IgG conjugated with horseradish peroxidase (anti-IgG F[ab]_2_ for F[ab]) (Jackson ImmunoResearch). IgM and IgA anti-OSP reactivity of plasma was measured using goat anti-human IgM and IgA conjugated with horseradish peroxidase (Jackson ImmunoResearch). Peroxidase activity was measured with the substrate 2,2-azinobis (ethylbenzthiazolinesulfonic acid) for plasma, o-phenylenediamine for purified IgG/M/A, and SuperSignal West Femto Maximum Sensitivity Substrate (ThermoScientific) for purified F[ab] fragments.

### Agglutination assay

Agglutination assays were performed as previously described with modification to match conditions used in assessing motility via high speed microscopy (22). Briefly, *V. cholerae* O1 classical Ogawa strain O395 was grown to mid-log phase in bovine heart infusion media. Bacteria were pelleted, washed twice with PBS, and diluted to OD_600_ of 0.1 with PBS. In a U-bottom microtiter plate blocked with BSA, 25 μl of bacterial samples was mixed with equal volume of day 7 patient sample to a final starting dilution/concentration of plasma 1:8; purified IgG 2.5, 1, or 0.5 μM depending on sample availability; purified IgG F[ab] 1 μM; or monoclonal antibody starting at concentration of 2.048 μM (320 mg/mL). The minimum agglutinating titer was determined by serial 2-fold dilutions of antibody sample. Plates were incubated at room temperature for 5 minutes and imaged using a UV imaging system (ChemiDoc, BioRad) to assess agglutination. If no agglutination was evident, the value was marked as > the most concentrated value.

### Viability assay

*V. cholerae* O1 classical Ogawa strain O395 was grown to mid-log phase in LB. Bacterial samples, adjusted to an OD_600_ of 0.1, were added to an equal volume of PBS or patient.s sample (heat-inactivated convalescent plasma or purified IgG for final antibody concentration of 1:256 for plasma and 0.25 μM and 2.5 μM for purified IgG). The mixture was incubated for 5 minutes at 37°C. The effect of antibody on bacterial viability was determined by serially diluting samples 10-fold and enumerating bacteria on LB plates. Experiments were performed in quadruplicate.

### Motility inhibition assay

*V. cholerae* motility was assessed using a modification of a previously reported approach (13). Briefly, *V. cholerae* O1 O395 were grown to mid-log-phase in LB broth and OD_600_ was adjusted to 0.1. This OD was found to approximate 3×10^6^ CFU/mL. Heat-inactivated convalescent antibodies were mixed with bacterial sample at a 1:1 dilution of acute or convalescent antibodies (final antibody concentration of plasma and purified IgG were at subagglutinating levels: 1:256 (plasma), 0.25 μM (polyclonal IgG); 0.5 μM (F[ab] generated from purified polyclonal IgG), and 10 μl of mixture was placed on a standard glass slide. Concentration dependence was also assessed for purified polyclonal IgG and monoclonal antibodies using 10-fold serial dilutions (2.5-0.025 μM for purified IgG; 2.5-0.00025 μM for monoclonal antibodies) using slides blocked with BSA. Motility inhibition of pooled samples of IgM and IgA were assessed at 0.25 μM; sample volumes were too low to assess motility individually for each patient using these antibody isotypes. PBS mixed 1:1 with bacteria was used as a negative control. Slides were incubated for 5 minutes at 37°C before visualization of motility by dark field microscopy (Nikon Eclipse Ti-E inverted microscope stand; Nikon Plan-Fluor 40x/0.75 objective; Nikon TI-DF Darkfield Condenser Dry 0.95-0.80; Hamamatsu ORCA-ER camera; MetaMorph imaging software). Images were taken at 400 msec exposure time over 100 frames with read out time frame 111.72 msec (movie play time of 1/30^th^ sec per frame). Frames 1, 50, and 98 were frozen as still shots and bacteria were counted as lines (motile bacteria) or dots (non-motile bacteria). Results were expressed as % motile (motile/total bacteria) and averaged across the three screen shots. To confirm that the antimotility effect was secondary to OSP-specific antibodies, we performed experiments with antibody samples that were first adsorbed with OSP:BSA. Antibody samples were first incubated overnight with OSP:BSA before mixing with the bacterial sample for microscopy. Adsorption was performed using a 1:100 molar ratio (antibody:OSP) for purified antibody, and 250 μg of OSP:BSA for diluted plasma samples. PBS mixed with OSP:BSA was used as a negative control. All assays were performed in at least triplicate.

### Electron microscopy

Scanning electron microscope (SEM) was performed at the Harvard University Center for Nanoscale Systems (CNS) using a FESEM Supra55VP microscope with a SE2 laser. *V. cholerae* O1 O395 was imaged in the presence or absence of anti-flagellin monoclonal antibody (AT11 0.025 μM), magnification range 45.3K-46.7K, or anti-OSP monoclonal antibody (G1 0.005 μM), magnification range 63.8K-66.5K.

### Neonatal challenge

The mouse neonatal *V. cholerae* challenge assay was used to assess protection afforded by OSP-specific antibodies as previously described (6). Briefly, three to five-day old CD-1 mice were separated from their dams for 3 hours and then orally inoculated with a 50 μL mixture containing 10^9^ *V. cholerae* O1 Inaba C6706 and PBS or anti-OSP monoclonal antibody (G1) at a final concentration of 0.25 μM. This strain was used due to the availability of mutant strains used in the competition assays (see below). Mice were then housed away from the dams at 30°C and monitored for survival for 30 hours. All mouse studies were approved by the MGH Institutional Animal Care and Use Committee.

### Competitive colonization assay

A single colony of each *V. cholerae* strain was inoculated into LB with Sm for C6706*lacZ*^*-*^ (referred to below as WT) and LB with Sm and Kan for C6706*lacZ*^*+*^ transposon mutants [rough (*VC0244*::Kan^r^) and non-motile (*motB*::Kan^r^)] and incubated at 37°C overnight on a roller drum. Cultures were then regrown to mid-log phase and the rough and comparator wild type (WT) strains were mixed together 1:1 (10^5^ organisms) and resuspended in 50 μL of LB without antibiotic mixed with PBS or 0.025 μM monoclonal antibody G1. A higher inoculum of the non-motile strain was required to enable sufficient recovery after infection for enumeration and calculation of a competitive index (input: 10^6^ mutant and 10^4^ comparator). Three to five-day old CD-1 infant mice were inoculated by oral gavage; and after 20 hours, the pups were sacrificed, and the proximal and distal thirds of the small intestines were isolated and mechanically homogenized in 5 mL of LB. Serial dilutions were plated on LB with Sm, X-gal and IPTG to enumerate bacteria and determine the input and output ratio of the comparator WT and competing rough or non-motile strain. The competitive index was calculated as the output ratio of competing strain/WT strain divided by the input ratio of competing strain/WT. The data represent infection of multiple litters on different days that were pooled for analysis. All mouse studies were approved by the MGH Institutional Animal Care and Use Committee.

### Statistical analysis

Results were expressed as medians and compared by a Wilcoxon matched-pairs signed rank sums test or Mann-Whitney U test for within group and between group comparisons as appropriate. Results of the survival curve analysis were compared by log rank testing. The threshold for statistical significance was a two-tailed p-value of <0.05. Prism 6 was used for all statistical analyses.

## ACKNOWLEDGMENTS

We would like to express our gratitude to the patients for participating in this study, and the field workers and research staff at the icddr,b, Bangladesh for their support and effort in sample collection and processing.

## Author Contributions

E.T.R., R.C.C., J.B.H, and F.Q. designed the study. F.C., A.I.K., T.R.B., and F.Q. assisted in enrolling and collecting patient samples for this analysis. R.C.C., M.K., J.M.T, M.H., K.I., A.A., L.MS., J.M.V., R.B., M.K. performed the experiments and analysis. J.B.H., J.W., A.R., R.C.K. generated and provided the human monoclonal antibodies. P.X. and P.K. produced *V. cholerae* OSP and OSP:BSA. M.K.W. produced *V. cholerae* C6706 strains. A.S.R., S.G., K.P.N, C.S.F. performed electron microscopy. R.C.C., M.K., J.M.T., M.H., K.I., A.A., J.M.V., D.T.L., R.C.L., A.W., F.C., A.I.K., S.B.C., T.R.B., J.B.H, F.Q. and E.T.R discussed the results. R.C.C, M.K., M.K.W., and E.T.R contributed to the writing. All authors edited and approved the final manuscript.

## SUPPLEMENTARY MATERIALS

**Table S1**. Cholera patient characteristics for polyclonal sera.

**Table S2**. *V. cholerae* strains used in this study.

**Table S3**. *V. cholerae* recovery in competitive colonization assays

**Fig. S1. Viability of *V. cholerae* is not affected by plasma or purified IgG in absence of complement**. Wild type *V. cholerae* classical strain O395 (OD 0.1) was mixed 1:1 with heat-inactivated convalescent plasma (diluted 1:256) or purified IgG (0.25 μM and 2.5 μM) and incubated for 5 minutes before serial dilutional plating on LB-agar. Each symbol represents the CFU/mL for a single experiment. Bar represents median with interquartile range. Differences between groups were assessed using a Mann-Whitney test. A p*-*value smaller than 0.05 was considered significant.

**Fig. S2. Inhibition of *V. cholerae* O1 O395 motility by pooled purified IgA and IgM plasma fractions of humans recovering from cholera in Bangladesh**. Pooled IgA (0.25 μM) and IgM (0.25 μM) factions of convalescent phase plasma of cholera patients (N=10) also block *V. cholerae* (OD 0.1) motility, and this effect is reduced by adsorption with *V. cholerae* OSP. d2:day 2; d7:day 7; d7 ads: day 7 adsorbed with OSP.

**Fig. S3. Scanning electron microscopy of *V. cholerae* mixed with monoclonal antibodies**. Scanning electron microscope (SEM) photos taken using a Supra55VP microscope with a SE2 laser. (A, B) *V. cholerae* O1 O395 (OD 0.1) and anti-flagellin monoclonal antibody (AT11 0.025 μM), magnification range 45.3K-46.7K. (C, D) *V. cholerae* O1 O395 (OD O.1) and anti-OSP monoclonal antibody (G1 0.005 μM), magnification range 63.8K-66.5K.

**Movie S1. *V. cholerae* motility after incubation with PBS**. High speed looped video microscopy after a 5-minute incubation of *V. cholerae* O1 O395 (OD_600_ 0.1) with phosphate-buffered saline. Note shooting star motility of individual *V. cholerae*.

**Movie S2. *V. cholerae* motility after incubation with anti-FlaA monoclonal antibody**. High speed looped video microscopy after a 5-minute incubation *V. cholerae* O1 O395 (OD_600_ 0.1) with a human monoclonal antibody targeting flagellin A (AT11; 0.25 μM). Note shooting star motility of individual *V. cholerae*.

**Movie S3. *V. cholerae* motility after incubation with anti-OSP monoclonal antibody**. High speed looped video microscopy after a 5-minute incubation *V. cholerae* O1 O395 (OD_600_ 0.1) with a human mAb targeting OSP (A4; 0.25 μM). Note significant diminution of shooting star motility of individual *V. cholerae* compared to what is observed following incubation with PBS (Movie S1) and anti-Flagellin specific monoclonal antibody (Movie S2).

